# Deacon: fast sequence filtering and contaminant depletion

**DOI:** 10.1101/2025.06.09.658732

**Authors:** Bede Constantinides, John Lees, Derrick W Crook

**Affiliations:** Nuffield Department of Medicine, University of Oxford, Oxford, UK; School of Biosciences, University of Birmingham, Birmingham, UK; European Molecular Biology Laboratory, European Bioinformatics Institute, Wellcome Genome Campus, Hinxton, UK; The National Institute for Health Research Oxford Biomedical Research Centre, University of Oxford, Oxford, UK

## Abstract

**Motivation:** Realising the value of large DNA sequence collections demands efficient search and extraction of sequences of interest. Search queries may vary in size from short gene sequences to multiple whole genomes that are too large to fit in computer memory. In microbial genomics, a routine search application involving both large queries and large collections is the removal of contaminating host genome sequences from microbial (meta)genomes. Where the host is human, *sensitive* classification and excision of host sequences is usually necessary to protect host genetic information. *Precise* classification is also critical in order to retain microbial sequences and permit accurate microbial genomic analysis. While human pangenomes have been shown to increase sensitivity of human sequence classification, existing bioinformatic host depletion approaches have either limited precision when used with metagenomes or large computing resource requirements.

**Results:** We present Deacon, an efficient and versatile sequence filter for raw sequence files and streams. We demonstrate its leading accuracy for the task of host depletion, using less computing resource than existing approaches. By querying a human pangenome index for minimizers contained in each input sequence, Deacon is able to accurately classify and discard diverse human sequences from long reads at over 250Mbp/s with a commodity laptop. We present validation of classification sensitivity, specificity and speed with simulated short and long reads for diverse catalogues of human, bacterial and viral genomes alongside existing methods. Beyond host depletion, Deacon is well suited to common sequence search and filtering applications, particularly those involving large queries. Capable of indexing a human genome in under 30s, Deacon is equipped to rapidly compose custom minimizer indexes using set operations, facilitating efficient search and filtering of massive sequence datasets using gigabase queries.

**Availability and implementation:** Deacon is implemented as an MIT-licensed command line tool written in Rust and packaged with Bioconda. Code is available from https://github.com/bede/deacon.

## 1 Introduction

The growing volume of generated and deposited DNA sequences demands ever more efficient methods for searching and filtering vast sequence collections. While search applications often involve short queries containing e.g. motifs or gene sequences, increasingly routine tasks such as microbial host decontamination involve probing large sequence collections with equally large queries comprising many whole genomes, posing a significant computational challenge. Thankfully, sequence sketching approaches using *k*-mers are well suited to problems of this nature, enabling compression of large genome collections to fit within computer memory. First conceived for document fingerprinting (Schleimer *et al*., 2003), *minimizer* sketching has seen particularly widespread adoption in bioinformatics, being effectively applied to problems ranging from genome alignment (Li, 2018) to *de novo* assembly (Ekim *et al*., 2021).

Here we describe a versatile tool for fast online searching and filtering of DNA sequences in files or streams using minimizer sketches. We demonstrate its accuracy and efficiency for an application motivating its development: classification and removal of host sequences from microbial sequencing reads, often called host depletion. Accuracy is of great importance in this context: sensitive classification and excision of host sequences is typically obligatory—albeit to an extent poorly defined—to protect the private genetic information of the host (Tomofuji *et al*., 2023). Yet precise classification and retention of non-host sequences is also critical for correct microbial genome and metagenome analysis. Speed and memory requirements are also concerns, particularly for rapid diagnostic applications (Lee and Pai, 2017; Street *et al*., 2025), and/or in remote locations without access to specialist computing resource (Quick *et al*., 2016).

Existing bioinformatic host depletion methods broadly rely on *i)* retention of sequences aligning to a target genome (Hunt *et al*., 2022), *ii)* depletion of sequences aligning to a host genome (Constantinides *et al*., 2023), *iii)* depletion of sequences containing host genome *k*-mers (Bushnell, 2014), *iv) k*-mer-based taxonomic classification (Hall and Coin, 2024), or *v)* combinations thereof (Rumbavicius *et al*., 2023; Seidel *et al*., 2025). While human pangenomes have previously been shown to increase the sensitivity of human sequence classification across diverse ancestries, we find that existing approaches exhibit either limited precision and/or require prohibitively large resources when used with human pangenomes. We demonstrate this by benchmarking sensitivity, specificity, balanced accuracy, speed and memory usage with simulated short and long reads for diverse catalogues of real human, bacterial and viral genomes.

## 2 Materials and Methods

Deacon is implemented as a standalone open source command line utility and library written in Rust. The index function constructs and saves a minimizer hash set from sequences contained in a target FASTA or FASTQ file. Prior to hashing, minimizers are canonicalised by selecting between each minimizer and its reverse complement. The resulting index can then be used for filtering unseen arbitrary sequence collections with the filter command, which accepts an index and a collection of query sequences as one or two (paired) input files in FASTA or FASTQ format, or else a standard input stream. During filtering, each distinct minimizer in each query sequence is canonicalised, hashed and tested for membership within the index hash set in memory. Canonical minimizer computation is accelerated using SIMD instructions (Groot Koerkamp and Martayan, 2025). Minimizers are hashed with the SIMD-accelerated XXH3 hash function. Match threshold is customisable as either a required integer number of minimizer hits per query, or a required proportion of minimizer hits per query. The user can specify whether to retain or discard matched queries according to use case. Gzip and Zstandard compression formats are handled automatically for input and output, and Deacon supports streaming input and output for single and paired sequences alike without temporary file creation.

### Benchmarking

Deacon 0.4.0’s performance for host contaminant filtering was evaluated alongside existing alignment-based (Langmead and Salzberg, 2012; Li, 2018) tool Hostile 2.0.0 (Constantinides *et al*., 2023) and *k*-mer-based (Wood *et al*., 2019) tools NoHuman 0.3.0 and Detaxizer 1.1.0 using default parameters (Hall and Coin, 2024; Seidel *et al*., 2025). Deacon’s panhuman-1 index was used, comprising minimizers from 47 diploid Human Pangenome Reference Consortium (HPRC) Year 1 assemblies, CHM13v2.0 and GrCh38.p14 (Liao *et al*., 2023; Nurk *et al*., 2022). Hostile’s default index comprises the human T2T-CHM13v2.0 reference and human leukocyte antigen (HLA) sequences, while NoHuman uses a Kraken2 index built from HPRC sequences, and Detaxizer uses the Kraken2 Standard-8 index by default in Kraken2 mode. Detaxizer was further evaluated using the full size 67GB Kraken2 Standard index dated 2025-04-02. During evaluation, the Detaxizer pipeline required modification to run to completion without memory errors on a Linux (x86_64) machine with 128GB of RAM using either index size. These modifications were contributed upstream as a merge request in the author’s public Git repository. We did not evaluate the approach proposed by Guccione et al. (2025) due to its pipeline requiring *>*1TB of RAM.

### Evaluation datasets

Simulated human, bacterial and viral genome catalogues were used to measure host classification sensitivity, specificity, and balanced accuracy. Short 2×150bp Illumina reads with 1% error were simulated with DWGSIM (Homer, 2022), while long ONT reads with approximately 4% error were simulated with PBSIM3 (Ono *et al*., 2022). Human reads were simulated for CHM13v2.0 and 12 HPRC assemblies each representing distinct human populations from Africa, Asia, Europe, and the Americas. Simulated bacterial reads comprised all 988 published complete FDA-ARGOS genomes (Sichtig *et al*., 2019), while simulated viral reads comprised all 17,900 complete NCBI RefSeq virus records as of 2025-02-25. Each constituent genome was simulated at 10x depth.

### Index design

Deacon’s default panhuman-1 index comprises the union of minimizer hashes (*k*=31, *w*=15) contained in the HPRC Year 1 assemblies for 47 samples (maternal and paternal haplotypes) and the human reference genomes GrCh38.p14 and T2T-CHM13v2.0 (Nurk *et al*., 2022). In order to slightly increase Deacon’s retention of microbial sequences, minimizers common to both the human pangenome and FDA-ARGOS bacteria or RefSeq virus genomes were removed using Deacon’s index subtraction functionality, resulting in removal of 20,741/409,935,039 (0.0051%) human pangenome minimizers.

## 3 Results

Henceforth, we refer to sensitivity as the proportion of removed human base pairs, and specificity as the proportion of retained microbial base pairs, with balanced accuracy representing the mean of both. We refer to 1-sensitivity as false negative rate (FNR), representing misclassified host sequences, and 1-specificity as false positive rate (FPR), representing misclassified microbial sequences. Results are summarised in Figure 1a. Figure 1b shows classification sensitivity for additional human genomes from 12 individuals with distinct ancestries.

**Fig. 1.**
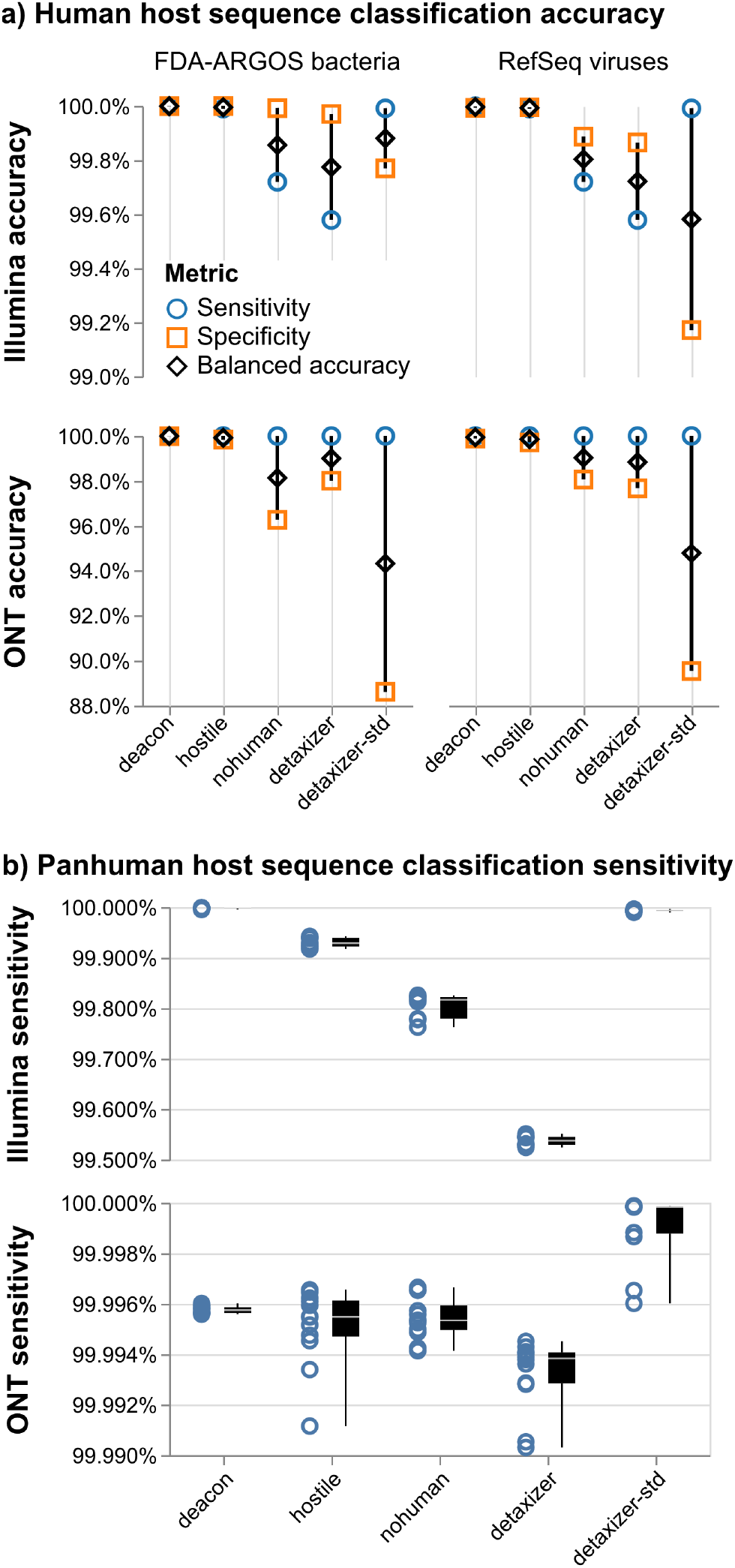
Host sequence classification accuracy. **a)**: Sensitivity, specificity and balanced accuracy measured using simulated short (top) and long (bottom) reads. Baseline sensitivity measured using human CHM13v2.0 reads. Specificity was measured using reads comprising 988 complete bacterial genomes (left) and 17,900 virus sequences (right) reflecting complex metagenomic backgrounds. **b)** Panhuman host sequence classification sensitivity for short (top) and long (bottom) reads simulated for 12 HPRC genome assemblies for individuals from Africa, Asia, Europe, and the Americas.

### Sensitivity of host sequence classification

For short read Illumina human pangenomes, Deacon was the most sensitive classifier with a median FNR of 1 *×* 10^−5^, followed by Detaxizer Standard (7 *×* 10^−5^), Hostile (7 *×* 10^−4^), NoHuman (2 *×* 10^−3^) and Detaxizer (5 *×* 10^−3^). For simulated long read ONT human pangenomes, Detaxizer Standard was the most sensitive with a median FNR of 2 *×* 10^−6^, followed by Deacon (4 *×* 10^−5^), Hostile (5 *×* 10^−5^), NoHuman (5 *×* 10^−5^) and Detaxizer (6 *×* 10^−5^). The FNR therefore varied between tools by one to two orders of magnitude.

### Specificity of host sequence classification

For short read Illumina bacterial genomes, Hostile was the most specific classifier with an FPR of (6 *×* 10^−7^), followed by Deacon (5 *×* 10^−6^), NoHuman (8 *×* 10^−5^), Detaxizer (3 *×* 10^−4^) and Detaxizer Standard (2 *×* 10^−3^). For short read Illumina viral genomes, the same ranking was observed, albeit with less difference between FPRs for Hostile (5 *×* 10^−5^) and Deacon (7 *×* 10^−5^). For long read ONT bacterial genomes, Deacon was the most specific with an FPR of 1 *×* 10^−4^, followed by Hostile (2 *×* 10^−3^), Detaxizer (2 *×* 10^−2^), NoHuman (4 *×* 10^−2^) and Detaxizer (1 *×* 10^−1^). For long read ONT viral genomes, Deacon was also the most specific (1 *×* 10^−3^), followed by Hostile (3 *×* 10^−3^), NoHuman, Detaxizer (2 *×* 10^−2^), and Detaxizer Standard (1 *×* 10^−1^). The FPR therefore varied between tools by three to four orders of magnitude.

### Balanced accuracy of host sequence classification

Deacon exhibited the highest balanced accuracy in all conditions followed closely by Hostile. Kraken2-based approaches NoHuman and Detaxizer exhibited lower balanced accuracy driven by relatively low specificity, with Detaxizer Standard consistently exhibiting the lowest balanced accuracy behind Nohuman And Detaxizer.

### Speed and memory usage

Speed and peak memory usage figures are shown in Tables 1, S1 and S2. Deacon was fastest at filtering both host and microbial long reads while using least memory, being 1.8-1.9x faster than Kraken2-based NoHuman, and 2.9x-53x faster than alignment-based Hostile. NoHuman was 1.1-1.3x faster than Deacon at processing short reads and 1.2x-20x faster than Hostile, with Hostile using the least memory. Hostile was an order of magnitude slower to process host reads than microbial reads. Neither Deacon nor Hostile created temporary files, while NoHuman decompressed its entire input to file during execution. The impact of input/output compression on Deacon’s performance was also examined (Table S1), during which long read decontamination speeds of up to 249Mbp/s (Zstandard compression) and 292Mbp/s (uncompressed) were recorded. Using Zstandard compression increased Deacon’s speed by between 28% and 44% over Gzip.

**Table 1.** Speed and memory use of host depletion measured on a laptop computer using simulated human genomes and bacterial metagenomes, reading and writing gzip-compressed FASTQ with default parameters and 8 threads.

| Dataset | Genomes | Base pairs | Speed (Mbp/s) |  |  | Peak RAM (GB) |  |  |
| --- | --- | --- | --- | --- | --- | --- | --- | --- |
|  |  |  | Deacon | Nohuman | Hostile | Deacon | Nohuman | Hostile |
| Human genomic long reads | 1 | 31,172,756,092 | **160.2** | 91.0 | 3.0 | **4.9** | 5.7 | 8.0 |
| Human genomic short reads | 1 | 31,172,755,800 | 60.5 | **80.3** | 3.0 | 4.5 | 5.2 | **3.7** |
| Bacterial metagenomic long reads | 988 | 41,207,981,833 | **78.1** | 41.3 | 26.6 | **4.9** | 6.0 | 7.2 |
| Bacterial metagenomic short reads | 988 | 41,207,903,700 | 29.0 | **31.0** | 24.5 | 4.5 | 5.2 | **3.3** |

*Note: Shown are median and maximum respective speed and peak memory use for three trials using a 2021 Apple M1 Pro (arm64) MacOS system. See Table S1 for speeds measured with different I/O compression formats.*

## 4 Discussion

Our findings indicate that Deacon’s minimizer screening implementation classifies and filters diverse host sequences from microbial sequences with greater speed and accuracy than existing approaches. While similarly accurate as tested, our approach avoids the overheads of scaling existing read aligners to human pangenomes: although pangenome graph aligners succinctly represent pangenomes and overcome the poor memory scaling of linear read aligners, their complexity makes them relatively slow for filtering tasks. Exact alignment performed by read aligners is also wasteful in the context of host depletion, making host-heavy sequences slow to decontaminate. Conversely, Deacon saves time when processing host sequences. Careful combined application of minimizer seeding with gapped alignment could be explored to increase filtering accuracy at the expense of some time and memory. Gapped alignment would likely prove advantageous when matching sequences with much higher error rates and/or divergence than considered here.

Both of the Kraken2-based approaches tested (NoHuman and Detaxizer) exhibited much lower specificity than other methods evaluated here and than described in previous literature. These methods misclassified between 2% and 11% of simulated long ONT reads, warranting further investigation. This can partly be explained by our use of more than 10,000 distinct bacterial and viral taxa to assess specificity, including those sharing many *k*-mers with the human genome. Recent host depletion studies selected no more than 30 microbial species for evaluating accuracy (Forbes *et al*., 2025; Seidel *et al*., 2025; Gao *et al*., 2025) which we consider insufficient representation for metagenomic applications. It should be noted that NoHuman was developed for Mycobacterial host depletion, for which it is well suited; the *M. tuberculosis* genome shares no 31-mers with the human genome (Hall, 2023). We hope that future evaluations of metagenomic host depletion methods will incorporate both diverse host genomes and diverse microbial genomes. The observed imprecision of Kraken2-based methods is nevertheless striking in light of Kraken2’s widespread use for filtering metagenomes. Detaxizer’s specificity for ONT reads reduced dramatically when used with the larger Standard index than its default 8GB index, albeit with greater sensitivity, consistent with the observations of (Seidel *et al*., 2025). Our results suggest that Deacon, Hostile and similar approaches cause little inadvertent damage to viral and bacterial sequence datasets. Furthermore, as more reference quality human genome sequences continue to be published, these can be efficiently incorporated into more complete panhuman Deacon indexes, improving classification and decontamination of real human genomes.

Having evaluated Deacon’s performance for host contaminant depletion, we wish to emphasise its versatility for filtering of single and paired sequence files and streams, leveraging the SIMD-accelerated minimizer implementation of Groot Koerkamp and Martayan (2025) and SIMD-accelerated XXH3 hashing. Deacon 0.4.0 decontaminated our uncompressed long read test datasets at over 200Mbp/s on a laptop, which could likely be increased through optimisation, particularly for short reads where our current implementation is suboptimal. Adding a Bloom prefilter to reduce memory requirements could also be explored in future work. Deacon 0.4.0 uses 64 bit hashes of minimizers, permitting memory-efficient use of long *k >* 32 minimizers. This does present a small risk of hash collisions and thus spurious matches. A slower collision-free version of Deacon was developed and tested to verify that hash collisions do not occur across our publicly deposited test data with the panhuman-1 index. This suggests that hash collisions are unlikely to be a practical concern unless using very large custom indexes.

## Data availability

Source code is available from https://github.com/bede/deacon. Simulated reads for CHM13v2.0 human, FDA-ARGOS bacteria (988 complete genomes) and RefSeq viruses (17,900 accessions) are publicly deposited with corresponding simulation parameters to facilitate reuse (Constantinides, 2025a,b,c). Deacon’s public Git repository also contains instructions for simulating pangenomes, alongside data underlying figures and tables.

