## Supplementary information for "Deacon: fast sequence filtering and contaminant depletion"

Up to date information on installing and using this software can be found in the GitHub repository (`https://github.com/bede/deacon`) together with raw data used to construct tables and figures.

Table S1: Deacon speed for host depletion on an Apple M4 (arm64) system running MacOS 15 with 10 threads, using the panhuman-1 index and combinations of simulated FASTA and FASTQ input/output formats with and without Gzip and Zstandard compression.

| | | | Speed (Mbp/s) | | | | | |
|---|---|---|---|---|---|---|---|---|
| | | | Uncompressed | | Zstandard | | Gzip | |
| Dataset | Samples | Base pairs | fasta | fastq | fasta | fastq | fasta | fastq |
| Human long reads | 1 | 31.2 Gbp | 292 | 266 | 249 | 230 | 191 | 179 |
| Bacterial long reads | 988 | 41.2 Gbp | 230 | 192 | 152 | 140 | 106 | 97 |

*Note: Speeds shown are medians of three trials*
